# Quantitative Cross-Linking/Mass Spectrometry Reveals Subtle Protein Conformational Changes

**DOI:** 10.1101/055418

**Authors:** Zhuo A. Chen, Lutz Fischer, Salman Tahir, Jimi-Carlo Bukowski-Wills, Paul N. Barlow, Juri Rappsilber

**Affiliations:** Wellcome Trust Centre for Cell Biology, Institute of Cell Biology, School of Biological Sciences, University of Edinburgh, Edinburgh EH9 3BF, UK; Schools of Chemistry and Biological Sciences, University of Edinburgh, Edinburgh EH9 3JJ, UK; Department of Bioanalytics, Institute of Biotechnology, Technische Universität Berlin, 13355 Berlin, Germany

## Abstract

We have developed quantitative cross-linking/mass spectrometry (QCLMS) to interrogate conformational rearrangements of proteins in solution. Our workflow was tested using a structurally well-described reference system, the human complement protein C3 and its activated cleavage product C3b. We found that small local conformational changes affect the yields of cross-linking residues that are near in space while larger conformational changes affect the detectability of cross-links. Distinguishing between minor and major changes required robust analysis based on replica analysis and a label-swapping procedure. By providing workflow, code of practice and a framework for semi-automated data processing, we lay the foundation for QCLMS as a tool to monitor the domain choreography that drives binary switching in many protein-protein interaction networks.

**Abbreviations:**

BS^3^: Bis[sulfosuccinimidyl] suberate
CLMS: Cross-linking/mass spectrometry
FDR: False discovery rate
HCD: Higher energy collision induced dissociation
LC-MS/MS: Liquid chromatography tandem mass spectrometry
LTQ: Linear trap quadrupole
MS2: Tandem mass spectrometry
QCLMS: Quantitative cross-linking/mass spectrometry
SCX: Strong cation exchange

## Introduction

Domain rearrangements of individual proteins act as molecular switches, govern the assembly of complexes and regulate the activity of networks. A typical example of a predominantly conformational change-driven finely tuned protein-protein interaction network is the mammalian complement system of ~40 plasma and cell-surface proteins. This is responsible for clearance of immune complexes from body fluids along with other hazards to health including bacteria and viruses. The study of such networks is hampered by the lack of tools to follow dynamic aspects of protein structures.

Cross-linking combined with mass spectrometry and database searching is a tool that can reveal the topology and other structural details of proteins and their complexes (1–3). It is currently unclear if dynamic information could also be obtained from such straight applications of cross-linking/mass spectrometry (CLMS) analysis. However, dynamic information could come from a comparison of the cross-links obtained from different protein states. We are developing a strategy to perform such a comparison in a rigorous manner by quantitative CLMS (QCLMS) (4) using isotope-labeled cross-linkers (5–8). In our strategy, distinct isotopomers of a cross-linker are used, to cross-link different stable conformers of the same, or very similar, protein molecules. The mass spectrometric signals of cross-linked peptides derived from different conformations can then be distinguished by their masses and thus quantified and related to conformational differences (Fig. 1A).

**Figure 1:**
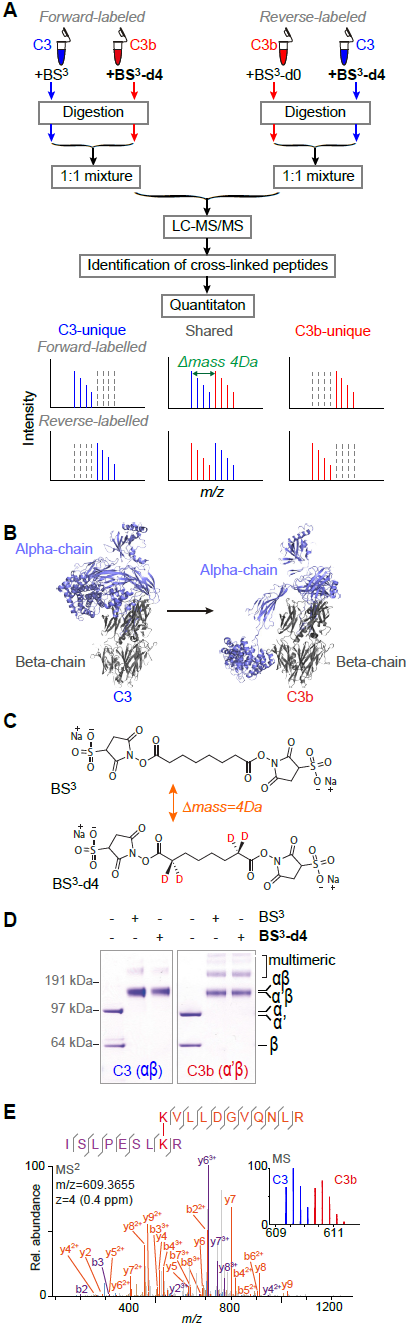
**Quantitative CLMS analysis of C3, C3b and iC3 in solution. (A)** The strategy of QCLMS using isotopologues of a cross-linker to compare two protein conformations. **(B)** The crystal structures of C3 (PDB|2A73) and C3b (PDB|2I07) with the α-chain (in C3)/α’-chain (in C3b) (blue) and the β-chains (grey) highlighted. **(C)** Chemical structure of cross-linkers BS^3^ and BS^3^-d4. **(D)** SDS-PAGE shows that BS^3^ (light cross-linker) and BS^3^-d4 (heavy cross-linker) cross-link C3 and C3b with roughly equivalent overall efficiencies, and that broadly similar sets of cross-linked protein products were obtained. **(E)** High-resolution fragmentation spectrum of BS^3^-cross-linked peptides ISLPESLK(cl)R-K(cl)VLLDGVQNLR that reveals a cross-link between Lys 267 and Lys 283. The mass spectrum of the precursor ion is shown (blue) in the inset; also present (red) is the signal of the precursor ion corresponding to the equivalent BS^3^-d4 cross-linked peptides.

Despite being straightforward in principle, quantifying cross-linked peptides is technically challenging. Currently available quantitative proteomics software is linked to protein identification workflows meaning that cross-link analysis is not routinely possible. Previously performed proof-of-principle work relied on an elementary computational tool, XiQ (4). A subsequent application (9) relied on manual data analysis, a subsequent protocol conspicuously left out the computational aspects (10). These works neglected to include replicated analysis and label-swaps; which are important standard procedures of quantitative proteomics. Recently, a workflow using replicated analysis and software mMass has been demonstrated using a calmodulin (17kDa) in presence and absence of Ca2+ (11). It remains to be seen if the approach scales to larger protein system. Additional experimental challenges derive from crosslinking itself. Cross-linking is known for low reproducibility. In any case, the signal intensity of an individual cross-linked peptide does not just depend on cross-linking efficiency. For example, additional variable chemical modifications of the peptides, such as methionine oxidation and reaction with the hydrolyzed cross-linker, may also depend on protein conformation. Furthermore, the network of cross-links that form within the molecule may influence the efficiency of protease digestion (11). Finally, peak interference or stochastic processes during data acquisition may prevent accurate quantitation. This is a general problem for quantification but it may especially affect cross-linking; cross-links are generally sub-stoichiometric and thus of low signal intensity in mass spectra.

To explore solutions to the challenges of QCLMS, we developed a workflow that involves replicated analysis, label-swap and some level of automation of the quantitation process, here utilizing the quantitative proteomics software tool Pinpoint (Thermo Scientific). This workflow has been successfully applied to study the conformational changes that involved in the maturation of the proteasome lid complex (12). We applied this workflow here to investigate key proteins in the complement system, a central player in human innate immune defenses. As the pivotal activation step of the complement system, C3 convertases excise the small anaphylatoxin domain (ANA) from the complement component C3 (184 kDa) leaving its activated form, C3b (175 kDa) (Fig. 1). Both C3 and C3b are stable, and comparisons of the crystal structures of both (C3 (13) and C3b (14)) (Fig. 1) revealed details of the structural rearrangements during this conversion. Using this comparison of C3 and C3b as a model system, we demonstrated and test in technical details the reliability of our workflow and usefulness of QCLMS. It is proved possible to infer, from our QCLMS data, the conformational changes that accompany cleavage of C3 to form C3b and then compare these to the difference between the two crystal structures. This both allowed cross-validation of the existing structures and revealed details of the relationship between cross-linking yields and conformational changes. Based on our experiences, we suggest a code of practice for the use of QCLMS in the study of protein dynamics.

## Experimental Procedures

### Protein preparation for cross-linking

Plasma-derived human C3 and C3b were purchased from Complement Technology, Inc. USA (and stored at −80 °C). Native C3 was depleted of low amounts of contaminating iC3 using cation-exchange chromatography (15). Chromatography was performed using a Mini S PC 3.2/3 column (GE Healthcare) at a flow-rate of 500 μl/min at 4 °C. Immediately after purification, C3 and C3b samples were exchanged, using 30-kDa molecular weight cutoff (MWCO) filters (Millipore), into cross-linking buffer (20 mM HEPES-KOH, pH 7.8, 20 mM NaCl and 5 mM MgCl_2_) with a final concentration of 2 ¼M. C3 and C3b samples used for “experiment I” and “experiment II” was prepared separately.

### Protein Cross-Linking

#### Experiment I

50 ¼g C3 and C3b were each, separately, cross-linked with either bis[sulfosuccinimidyl] suberate (BS^3^) (Thermo Scientific) or its deuterated analogue bis[sulfosuccinimidyl] 2,2,7,7-suberate-d4 (BS^3^−d4) (Fig. 1C), at 1:3 protein to cross-linker mass ratios, giving rise to four different protein-cross-linker combinations: C3+BS^3^, C^3^+BS3−d4, C3b+BS^3^ and C3b+BS^3^−d4. After incubation (two hours) on ice, reactions were quenched with 5 μl 2.5 M ammonium bicarbonate for 45 minutes on ice. For the monitoring of cross-linked products, an aliquot containing 5 pmol of cross-linked protein from each of the above four reaction mixtures were subjected to SDS-PAGE using a NuPAGE 4–12% Bis-Tris gel and MOPS running buffer (Thermo Fisher Scientific) according to the manufacturer’s instructions. The protein bands were visualized using the Colloidal Blue Staining Kit (Thermo Fisher Scientific) according to the manufacturer’s instructions.

Cross-linking reactions of C3 and C3b samples were repeated for “experiment II” as described for “experiment I”.

### Sample Preparation for Mass Spectrometric Analysis

#### Experiment I

Native C3 and C3b molecules each contain two, disulfide-linked, polypeptide chains. The monomeric (two-polypeptide chain) products of cross-linked C3 and C3b were resolved as distinct bands using SDS-PAGE. The products were in-gel reduced and alkylated, then digested using trypsin following a standard protocol (16). For quantitation, equimolar quantities of the tryptic products from the four cross-linked protein samples were mixed pair-wise, yielding two combinations: C3+BS^3^ and C3b+BS^3^−d4 (named here as forward-labeled); C3+BS^3^−d4 and C3b+BS^3^ (named here as reverse-labeled).

From each of these two samples, a 20–μg (40 μL) aliquot was taken and fractionated using SCX-Stage-Tips (17) with a small variation of the protocol previously described for linear peptides (18). In short, peptide mixtures were first loaded on a SCX-Stage-Tip in loading buffer (0.5% v/v acetic acid, 20% v/v acetonitrile, 50 mM ammonium acetate). The bound peptides were eluted in two steps, with buffers containing 100 mM ammonium acetate and 500 mM ammonium acetate, producing two fractions. These peptide fractions were desalted using C18-StageTips (19) prior to mass spectrometric analysis.

#### Experiment II

Preparation of two quantitation samples C3+BS^3^/C3b+BS^3^−d4 (forward-labeled) and C3+BS^3^− d4/ C3b+BS^3^ (reverse-labeled) were repeated as described for “experiment I”. A 4-μg (10 μL) aliquot of each sample was desalted using C18-Stage-Tips for mass spectrometric analysis without pre-fractionation.

### Mass Spectrometric Analysis

#### Experiment I

SCX-Stage-Tip fractions were analyzed using a hybrid linear ion trap-Orbitrap mass spectrometer (LTQ-Orbitrap Velos, Thermo Fisher Scientific) applying a “high-high” acquisition strategy. Peptides were separated on an analytical column that was packed with C18 material (ReproSil-Pur C18-AQ 3 μm; Dr. Maisch GmbH, Ammerbuch-Entringen, Germany) in a spray emitter (75–μm inner diameter, 8–μm opening, 250–mm length; New Objectives, Woburn, MA, USA) (20). Mobile phase A consisted of water and 0.5% v/v acetic acid. Mobile phase B consisted of acetonitrile and 0.5% v/v acetic acid. Peptides were loaded at a flow-rate of 0.7 μl/min and eluted at 0.3 μl/min using a linear gradient going from 3% mobile phase B to 35% mobile phase B over 130 minutes, followed by a linear increase from 35% to 80% mobile phase B in five minutes. The eluted peptides were directly introduced into the mass spectrometer. MS data were acquired in the data-dependent mode. For each acquisition cycle, the mass spectrum was recorded in the Orbitrap with a resolution of 100,000. The eight most intense ions with a precursor charge state 3+ or greater were fragmented in the linear ion trap by collision-induced disassociation (CID). The fragmentation spectra were then recorded in the Orbitrap at a resolution of 7,500. Dynamic exclusion was enabled with single repeat count and 60-second exclusion duration.

#### Experiment II

Non-fractionated peptide samples were analyzed using a hybrid quadrupole-Orbitrap mass spectrometer (Q Exactive, Thermo Fisher Scientific). Peptides were separated on a reversed-phase analytical column of the same type as described above. Mobile phase A consisted of water and 0.1% v/v formic acid. Mobile phase B consisted of 80% v/v acetonitrile and 0.1% v/v formic acid. Peptides were loaded at a flow rate of 0.5 μl/min and eluted at 0.2 μl/min. The separation gradient consisted of a linear increase from 2% mobile phase B to 40% mobile phase B in 169 minutes and a subsequent linear increase to 95% B over 11 minutes. Eluted peptides were directly sprayed into the Q Exactive mass spectrometer. MS data were acquired in the data-dependent mode. For each acquisition cycle, the MS spectrum was recorded in the Orbitrap at 70,000 resolution. The ten most intense ions in the MS spectrum, with a precursor change state 3+ or greater, were fragmented by Higher Energy Collision Induced Dissociation (HCD). The fragmentation spectra were thus recorded in the Orbitrap at 35,000 resolution. Dynamic exclusion was enabled, with single-repeat count and a 60 second exclusion duration.

### Identification of cross-linked peptides

The raw mass spectrometric data files were processed into peak lists using MaxQuant version 1.2.2.5 (21) with default parameters, except that “Top MS/MS Peaks per 100 Da” was set to 20. The peak lists were searched against C3 and decoy C3 sequences using Xi software (ERI, Edinburgh) for identification of cross-linked peptides. Search parameters were as follows: MS accuracy, 6 ppm; MS2 accuracy, 20 ppm; enzyme, trypsin; specificity, fully tryptic; allowed number of missed cleavages, four; cross-linker, BS^3^/BS^3^-d4; fixed modifications, carbamidomethylation on cysteine; variable modifications, oxidation on methionine, modifications by BS^3^/BS^3^–d4 that are hydrolyzed or amidated on the other end. The reaction specificity for BS^3^ was assumed to be for lysine, serine, threonine, tyrosine and protein N-termini. Identified candidates for cross-linked peptides were validated manually in Xi, after applying an estimated FDR of 3% for cross-linked peptides (Fischer and Rappsilber, submitted). We used 3%FDR as this was the best FDR that returned reasonable number of decoys to provide a meaningful FDR. List of identified cross-linked residue pairs were summarized based these cross-linked peptides. We also included additional cross-linked peptides that support identified cross-linked residue pairs for quantitation. These cross-linked peptides were identified in separate mass spectrometry runs with less than 3% FDR and manually verified spectra. The identification of these cross-linked peptides were transferred into the quantitation runs using “match between runs” based on high m/z accuracy and reproducible chromatographic retention time for MS1 signals. In addition, transferred identification were further verified in the quantitation runs with their MS signal pattern (either shown as doublet signals or singlet signals with 4D mass shift between paired label-swapped replicas). Identification information of all quantified cross-linked peptides and the annotated best-matched MS2 spectra for quantified cross-linked peptides are provided in Supplemental Table S1 and Supplemental File S1.

### Quantitation of cross-link data using Pinpoint software

Identified cross-linked peptides were quantified based on their MS signals. The quantitative proteomics software tool Pinpoint (Thermo Fisher Scientific) was used to retrieve intensities of both light and heavy signals for each cross-linked peptides in an automated manner. An input library for cross-linked peptides was constructed according to “General Spectral Library Format for Pinpoint Comma Separated Values” (Thermo Fisher Scientific) (Supplemental File S1). In the input library, the sequence of every cross-linked peptide was converted into a linear version with identical mass (Fig. 2A), following an idea from Maiolica et al. (13). Six modifications that were not listed in Pinpoint modification list were defined in “customized modifications”:

[BS3-OH]; mass, 156.07860; Site modified, KSTY
[BS3-NH2]; mass, 155.09460; Site modified, KSTY
[BS3d4-OH]; mass, 160.10374; Site modified, KSTY
[BS3d4-NH2]; mass, 159.11972; Site modified, KSTY
[Xlink]; mass, 27.98368; Site modified, K
[Xlinkd4]; mass, 32.00878; Site modified, K

**Figure 2:**
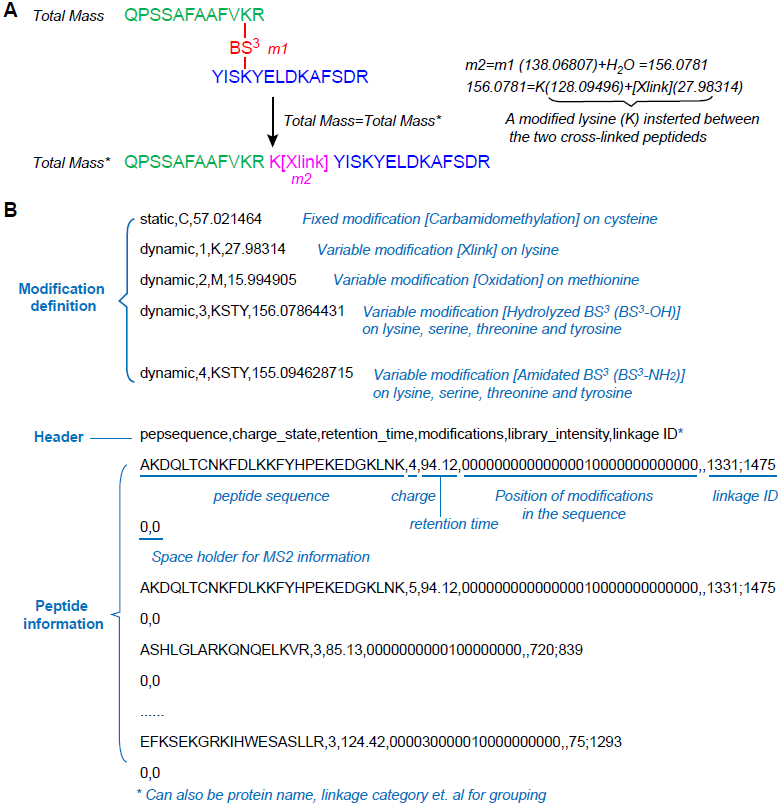
**Construction of Pinpoint peptide library for cross-linked peptides. (A)** The principle of converting a cross-linked peptide pair into a “linearized” format that has an identical mass. **(B)** An example Pinpoint-input peptide library for cross-linked peptides with its key contents annotated.
Figure 3: Quantitation of cross-linked peptides.

The three most abundant of the first four signals in the isotope envelope were used for quantitation. The error tolerance for precursor m/z was set to 6 ppm. Signals are only accepted within a window of retention time (defied in spectral library) ±10 minutes. Manual inspection was carried out to ensure the correct isolation of elution peaks. For each cross-linked peptide, elution peak areas of the light and the heavy signals were measured and reported as log2 (C3b/C3). “Match between runs” (22) was carried out for all cross-linked peptides in Pinpoint interface manually, based on high mass accuracy and reproducible LC retention time, hence quantitation was conducted for each identified cross-linked peptides in both forward-labeled and reverse-labeled samples even when they were only identified in one of them. In “experiment I”, if a cross-linked peptide was quantified in two SCX fractions, the average of fold-change (log2(C3b/C3)) in both fractions was reported. Within each of four biological (cross-link) replicas (experiment I forward-labeled, experiment I reverse-labeled, experiment II forward-labeled and experiment II revers-labeled), signal fold-changes of all quantified cross-linked peptides were normalized against their median, in order to correct systematic error introduced by minor shift on mixing ratio during sample preparation. C3b/C3 signal ratios of all quantified peptides in two experiments (four cross-link replicas) were listed in Supplemental Table S1.

The quantitation data was subsequently summarized at the level of unique residue pairs (cross-links). A cross-link was defined as a unique cross-link in either C3 or C3b only if all its supporting cross-linked peptides were observed as the corresponding singlet signals. Otherwise, cross-links were regarded as having being observed in both conformations. For a cross-link shared by C3 and C3b, the signal fold-change was defined as the median of all its supporting cross-linked peptides. Only those cross-links that were consistently quantified in both paired replica analyses (with label-swap) were accepted for subsequent structural analysis. Singlet cross-links were further confirmed by a mass shift of 4 Da, resulting from the label-swap. For cross-links observed as a doublet, the average of signal fold-changes from label-swapping replicated analyses was reported (Fig 3C). When a cross-link was quantified in both “experiment I” and “experiment II”, its fold-change in the two analyses was averaged and reported. All quantified cross-links are listed in Supplemental Table S2. Cross-links that were significantly enriched in C3 or C3b were determined using “Significance A” test from the standard proteomics data analysis tool Perseus (version 1.4.1.2) (21) based on log2(C3b/C3) values. The following parameters were used in Perseus for the test: “Side”: both; “Use for truncation”: P value; “Threshold value”: 0.05.

**Figure 3:**
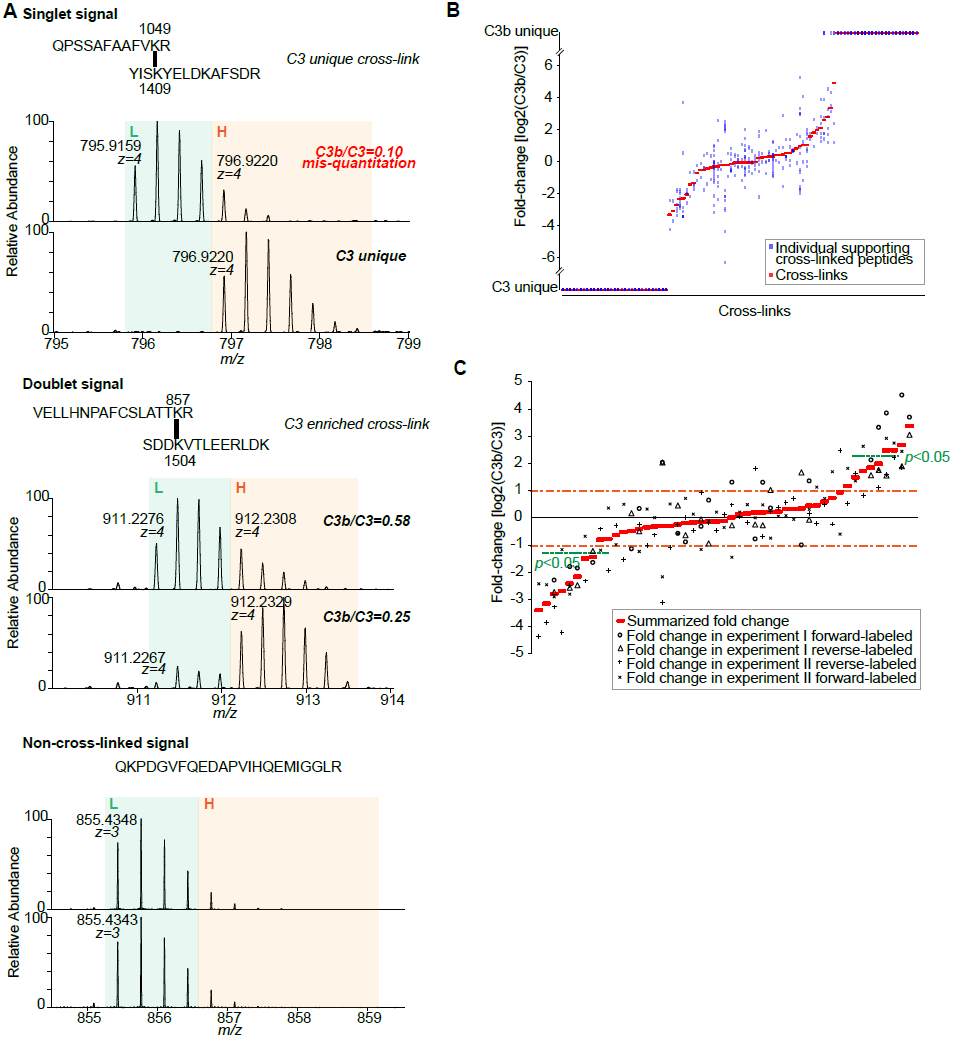
**Quantitation of cross-linked peptides. (A)** MS signals from paired quantitation analyses with label-swapping. These allow a clear distinction to be made between cross-linked peptides with a singlet signal, cross-linked peptides with a doublet signal, and a non-cross-linked peptide; these are all potentially similar and could therefore be confused in a non-paired quantitative analysis **(B)** Spread of quantitation data at the level of cross-linked peptides. For each of 104 quantified cross-links, the ratio of C3b to C3-derived signal is shown in red, while the ratio of C3b to C3-derived cross-link signal for its supporting, cross-linked peptides is plotted in blue **(C)** Ratios of C3b to C3-derived signals for 49 cross-links that were observed in both C3 and C3b. The summarized C3b/C3 ratios and C3b/C3 ratios observed in biological (cross-link) replicas are shown for each cross-linked residue pair. Cross-links that are significantly enriched are indicated (“Significant A” test *p*< 0.05 (21)).

### Comparison with crystal structures

To compare cross-linking data with X-ray crystallographic data, cross-links were displayed using PyMol (version 1.2b5) (23) in the crystal structures of C3 (PDB|2A73) and C3b (PDB|2I07). Cross-links were represented as strokes between the C-α atoms of linked residues. In the case of a residue missing from the crystal structures, the nearest residue in the sequence was used for display purposes. The distance of a cross-linked residue pair in the crystal structures was measured between the C–α atoms. Measured distances of linked residue pairs in crystal structures were compared to a theoretical cross-linking limit, which was calculated as side-chain length of cross-linked residues plus the spacer length of the cross-linker. An additional 2 Å was added for each residue as allowance for residue displacement in crystal structures. The following side-chain lengths were used for the calculation: 6.0 Å for lysine, 2.4 Å for threonine, 2.4 Å for serine and 6.5 Å for tyrosine. For example, for a lysine-lysine cross-link, the theoretical cross-linking limit is 27.4 Å. Solvent accessibility of cross-linked residues in the crystal structures were obtained using ‘Protein interfaces, surfaces and assemblies' service PISA at the European Bioinformatics Institute. (www.ebi.ac.uk/pdbe/protint/pistart.html) (24).

### Accession codes

The mass spectrometry proteomics data have been deposited to the ProteomeXchange Consortium (25) (http://proteomecentral.proteomexchange.org) via the PRIDE partner repository with the dataset identifier PXD001675.

## Results

### Quantitative cross-linking/mass spectrometry (QCLMS) of C3 and C3b in solution

To assess the abilities of QCLMS to reflect conformational changes, we studied the structurally well-characterized differences between C3 (PDB|2A73) and its activated cleavage product C3b (PDB|2I07). We cross-linked both purified proteins in solution using bis[sulfosuccinimidyl] suberate (BS^3^) and its deuterated analogue BS^3^-d4 (5–7) (Fig 1C) in four distinct protein-cross-linker combinations, generating C3+BS^3^, C3+BS^3^−d4, C3b+BS^3^ and C3b+BS^3^−d4. BS^3^ is a homo-bifunctional cross-linker containing an amine-reactive *N*-hydroxysulfosuccinimide (NHS) ester at each end. It reacts primarily with the ε-amino groups of lysine residues and the amino-terminus of a protein, however it can also react with hydroxyl groups of serine, tyrosine and threonine residues (26). Native C3 and C3b are each composed of two polypeptides chains (β and α for C3, p and α’ for C3b) connected by a disulfide bond. In all four cross-linking reactions, the two polypeptide chains of C3/C3b were efficiently cross-linked by BS^3^ or BS^3^-d4, and the resultant two-chain products of cross-linking could be isolated by SDS-PAGE (Fig. 1D) and subjected to in-gel trypsin digestion (16). A 1:1 mixture of C3+BS^3^ and C3b+BS^3^−d4 digests along with a “label-swapped” replica (C3+BS^3^−d4 and C3b+BS^3^ digests) were analyzed by liquid chromatography-tandem mass spectrometry (LC-MS/MS) (27). Cross-linked peptides were identified by database searching and then quantified based on their MS signals (Fig. 1E).

Wherever C3 and C3b have regions of identical structure, cross-linked peptides should be seen in mass spectra as 1:1 doublet signals (separated by 4 Da). In contrast, where C3 and C3b structures differ, unique cross-links may occur and these will lead to singlets in mass spectra (Fig. 1A). Signal interferences, extended isotope envelopes or experimental variations may affect quantification. Experimental variations are addressed by replica analysis. Swapping the use of labels in the replica addresses problems arising from extended isotope envelopes and signal interference. Importantly, the label-swap enabled quantitation of cross-linked peptides with singlet signals by generating signature signal patterns (Fig 1A); it also prevented singlet signals of BS^3^ cross-linked (light) peptides from being mistakenly quantified as doublets when the heavy and the light signals of cross-linked peptides overlap (Fig. 3A). Swapping the use of labels also assists in the identification of cross-links. Singlet signals move by 4 Da, while doublet signals remain as doublets. This behavior of signals corresponding to cross-linked peptides is also clearly distinct from those of peptides contain no cross-linked amino acid residues (Fig. 3A).

In experiment I, we conducted strong cation exchange (SCX)-based fractionation to increase the number of identifiable cross-linked peptides (Fig. 4A). Interestingly, only 20% of all unique cross-linked peptide pairs (25/127) eluted in more than a single fraction. 24 of these 25 cross-linked peptides eluted in two fractions in both forward-labeled and reverse-labeled samples (one only quantified in the reverse-labeled) showing good reproducibility of SCX fractionation. For 49 quantified events of these 25 cross-linked peptides that were identified and quantified in two sequential SCX fractions, we observed that the C3b/C3 signal ratios were highly reproducible (100% for C3/C3b unique cross-linked peptides and *R*^2^=0.95 for cross-linked peptides quantified with ratios, Fig. 4B). We conclude that SCX appears to be compatible with QCLMS and a worthwhile step in the procedure while cross-linked peptides can be quantified with high technical reproducibility.

**Figure 4:**
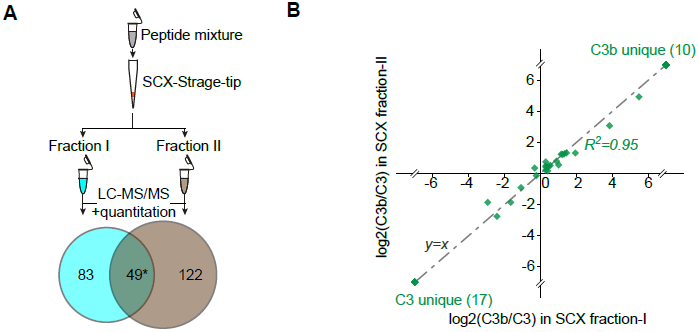
**Strong cation exchange based enrichment for cross-linked peptides is compatible with QCLMS analysis**. **(A)** Strong cation exchange (SCX)-based enrichment using SCX-Stage Tips applied in QCLMS analysis of C3 and C3b. **(B)** Reproducibility of C3b/C3 signal ratios of 49 quantified events of 25 unique cross-linked peptides in two SCX fractions in which they were detected.

In an ideal case, the ratio of the component signals of a doublet would simply be a function of the relative likelihood of linking a particular residue pair in each of the two different protein conformations. As shown above, good agreement on the signal ratio of identical crosslinked peptides quantified in sequential SCX fractions was observed in both forward-labeled and reverse-labeled experiments in experiment I, suggesting high technical reproducibility of our analysis. However, we observed that different cross-linked peptide pairs that contained the same pair of cross-linked residues, did not share the same component signal ratios (Fig. 3B). For example, within a single experiment, for the six different cross-linked peptides that covered the residue pair Lys1 and Lys646, the (C3b/C3) signal ratio varied between 0.3 and 1.9 (Supplemental Table S1). This by far exceeds the variation seen in technical replicates. They resulted from methionine oxidation and also missed cleavages and thus may link to sample preparation or to the impact of conformation on proteolysis cleavage efficiencies. To translate data on multiple peptide pairs containing the same cross-linked residues into a single data point for that residue pair, we took the median ratio [i.e. log2(C3b/C3)] of all supporting cross-linked peptide pairs for each residue pair. This is equivalent to the procedure used for the inference of protein ratios from peptide ratios in standard quantitative proteomics.

### Semi-automated quantitation for cross-linked peptides using Pinpoint

A practical challenge for quantifying cross-link data is how to capture efficiently the signal intensities of cross-linked peptides from a large quantity of mass spectrometric raw data. Available quantitative proteomics software does not accommodate cross-linked peptides (4). A previous protocol for QCLMS resorted to manual quantification (10). Manual quantitation relies more on user expertise and subjective criteria; in respect to automated analysis, it is less comparable between labs. When dataset size increases, manual quantitation becomes increasingly time consuming and impractical. Furthermore, any repetitive task done by a human is error-prone. To alleviate this we developed a semi-automated workflow using the quantitative proteomics software package, Pinpoint. The resultant workflow utilizes the well-established functionality of Pinpoint to retrieve intensities of both light and heavy version of every crosslinked peptide in an automatic manner. Furthermore, the user interface of Pinpoint provides, when necessary, a platform for visualizing and validating the quantitation results of any chosen cross-linked peptides. Hence, improvements to the accuracy of quantitation by introducing knowledge-based expertise can be achieved easily and rapidly.

Pinpoint is normally restricted to work with single peptides. Therefore it was necessary to convert our cross-linked peptide pairs into a “linearized” single-peptide format, following a previous approach developed for database searches (16) (Fig. 2A). This allowed generation of a tailored input library of cross-linked peptides (Fig. 2B) based on “General Spectral Library Format for Pinpoint” (Supplemental File S1). In general, Pinpoint requires a very basic set of information to quantify a cross-linked peptide: amino acid sequence, precursor charge state, retention time, and chemical modifications. A consequence of this minimal requirement of information is that quantification can be performed in Pinpoint independently of the method used for identifying the cross-linked peptides in the first place.

Pinpoint calculates the theoretical m/z of each converted (“linearized”) cross-linked peptide and identifies its signal (within 6 ppm error tolerance in our case) in the raw MS data. Although isotope-related shifts in retention time often occur, in most cases, the retention time of the BS^3^ and BS^3^-d4 cross-linked versions of a cross-linked peptide in LC-MS/MS overlap to some extent. In order to accurately define singlet and doublet signals of cross-linked peptides, we programmed Pinpoint to retrieve intensity information of both light and heavy signals for each cross-linked peptide by including both a BS^3^ cross-linked and a BS^3^-d4 cross-linked version of them in the input library.

As discussed above, accurate quantitation of a cross-linked peptide relies on consistent read-out from both replicates of the label-swap analysis. This is especially important for crosslinked peptides that are unique in one of the two conformations. However, cross-linked peptides were not always fragmented and identified in each replica. Such “under sampling” of signals is common in shotgun proteomics and likely exasperated due to the generally low abundance of cross-linked peptides. The Pinpoint interface also provides a platform to conduct “Match between runs” (22). The high mass accuracy achieved in the analysis of the high-resolution Orbitrap data and reproducible LC retention time facilitated this transfer of peptide identities. Thus, all identified cross-linked peptides were quantified in both replica experiments even though they were not necessarily identified in both replicas.

The aggregated intensity of each cross-linked peptide was calculated from the summed area of the three most intense peaks among the first four peaks in the isotope envelope. In cases where a cross-linked peptide was identified with more than one charge state, intensities derived from the different charge states were automatically combined by Pinpoint. Pinpoint did not select correctly the start or end of an elution peak in every instance. Manual curation was therefore still necessary, albeit largely supported by the Pinpoint interface. In addition, crosslinked peptides were discarded for quantitation if they did not have proper signals for the first three peaks of their isotope envelopes.

Consequently, for each cross-linked peptide, Pinpoint provided intensities for both light (BS^3^) and heavy (BS^3^-d4) signals. The results were exported into a.csv file. The subsequent processes for generating the final quantitation results were carried out using a Microsoft Excel spreadsheet. This included calculating the C3b/C3 signal ratio for each quantified event of cross-linked peptides, normalizing C3b/C3 signal ratio within each cross-link replica, summarizing C3b/C3 signal ratios for each cross-linked residue pair from the C3b/C3 signal ratios of its supporting cross-linked peptides; and then combining the outcomes of quantitation for the label-swapped replicas and for two experiments (experiment I and experiment II).

### Cross-linking confirms in solution, C3 to C3b conformational transition

In total, we quantified 104 unique cross-linked residue pairs (cross-links) that could be divided into three groups based on their MS1 signal types: 31 C3-unique cross-links, 24 C3b-unique cross-links and 49 cross-links observed in both proteins. When the “Significance A” test from the standard proteomics data analysis tool Perseus (version 1.4.1.2) (21) was applied to the 49 cross-links observed in both C3 and C3b, based on their log2(C3b/C3) values, three subgroups appeared: 37 showed no significant change (named here mutual), four linked residue pairs were significantly enriched (*p*<0.05) in C3b and eight linked residue pairs were significantly enriched in C3 (*p*<0.05). The conformational differences/similarities between C3 and C3b were revealed by the locations of these 104 quantified cross-links (Fig. 5A). 31 C3-unique cross-links and 24 C3b-unique cross-links highlighted where the structures of C3 and C3b differs; 37 cross-links that showed no significant change would suggest structural features that are unaffected by cleavage of C3 to C3b; eight C3-enriched and four C3-enriched cross-links potentially reflected minor conformational changes. These quantified groups and subgroups were not randomly distributed, neither in the primary sequences nor in the 3D structures of C3 (PDB|2A73) (13) and C3b (PDB|2I07) (14) (Figs. 5B, C, D, E). The structural differences and similarities between C3 and C3b as deduced based on our QCLMS data are in agreement with the crystal structures of the two proteins.

**Figure 5:**
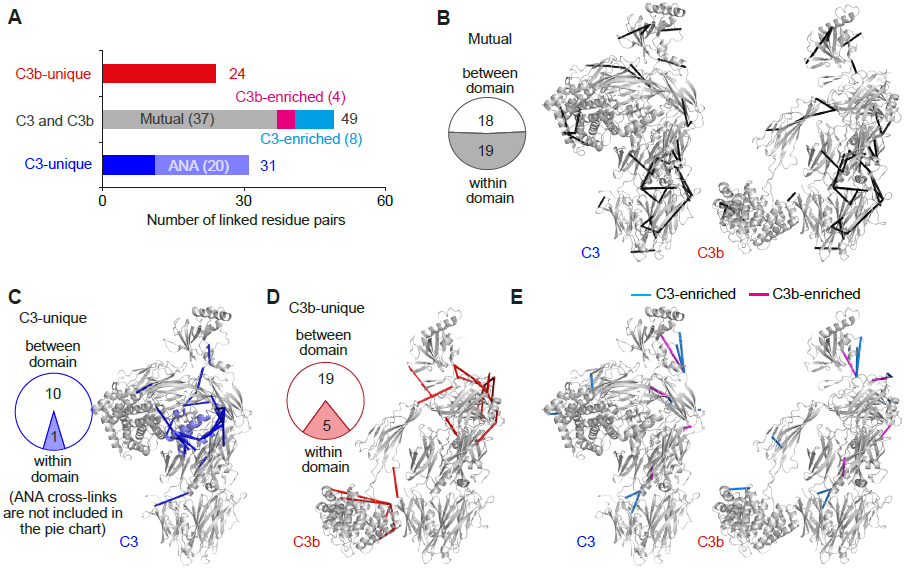
**Quantitative CLMS analysis reveals the domain rearrangements, in solution, which accompany the activation of C3. (A)** 104 unique cross-links that were quantified in C3 and C3b samples were allocated to three groups: 31 are unique to C3 (20 involve the ANA domain); 24 are unique to C3b; 49 were observed in both C3 and C3b (of these, four are statistically enriched in C3b, eight are enriched in C3 and 37 (denoted “mutual”) are equally represented in both C3 and C3b. **(B)** The 37 mutual cross-links (black) are displayed on the crystal structures of C3 (PDB|2A73) and C3b (PDB|2I07). Of these, 19 are intra-domain, consistent with the conserved structures of these domains in C3 versus C3b. The remaining (18) links pairs of domains whose relative positions are similar in C3 and C3b. **(C)** As displayed in the C3 structure, C3-unique cross-links (blue) lie mainly within the C3-specific ANA domain (light blue) and between domains in the α-chain, which are significantly rearranged upon formation of C3b. **(D)** The C3b-unique cross-links (red) occur primarily between domains within the rearranged α-chain and at the new interface formed between the TED and MG1 domains (displayed on PDB|2I07).

Conversion from C3 to C3b is triggered by proteolytic cleavage of the 7-kDa anaphylatoxin (ANA) domain from the N-terminus of the alpha chain. It is therefore not surprising that 20 out of the 31 residue pairs that were found to be cross-linked only in C3 involved ANA. Six of these 20 residue pairs included ANA residues as one partner and residues of MG3, MG8 and TED as the other. Thus these C3-exclusive ANA-specific cross-links clearly define a spatial location for ANA in the C3 molecule that is wholly consistent with the crystal structure (Fig. 5C). The remaining 84 cross-links (11 unique to C3, 24 unique to C3b, and 49 mutual ones) (Figs. 5B, C, D E), report on the extent and nature of rearrangements of the 12 non-ANA domains, namely C345C, TED, CUB, LNK and eight MG domains.

The data confirmed that the spatial arrangement of the domains of the β-chain is conserved following cleavage of the alpha chain: only two cross-links unique to either C3 or C3b involved residues in this chain, compared to 33 C3-unique or C3b-unique cross-links for α-chain domains. The cross-links in the β-chain that are conserved between C3 and C3b occur within (13 cross-links) or between (10 cross-links) its seven intact domains (six MG domains and the LNK domain). In contrast, the remaining domains of the α-chain appears to rearrange extensively following ANA excision: A total of 28 of the 33 C3-unique or C3b-unique cross-links in the α-chain are between domains and only five are within domains; this compares to the identification of nine out of the 12 preserved (not unique to C3 or C3b) α-chain cross-links within domains. In essence, all but one of the domains remain largely unaltered in structure following activation of C3, despite their arrangement changing. An exception is a set of four C3b-unique cross-links identified within MG8. This suggests that conformational changes occur within MG8, to a greater extent than in the other MG domains, following C3 cleavage

Inspection of pairs of amino acid residues that were cross-linked exclusively either in C3 or in C3b illuminates three major rearrangements in the α-chain that accompany the conversion of C3 to C3b. These observations, made in solution, strongly agree with the conformational changes inferred from comparing the crystal structures of C3 (PDB 2A73) and C3b (PDB 2I07) (Fig. 5C, D; Supplemental Table S2). First, residues within the α’-N-terminal (NT) segment (the region of the C3b α-chain that becomes the new terminus after ANA cleavage) are involved in four C3-unique cross-links and six C3b-unique cross-links. These cross-links captured, in C3, the proximity of the α-NT segment to ANA and MG8, In C3b the cross-links confirmed relocation of α’-NT from the MG8/MG3 side to the opposite, C345C/MG7, side of the structure (14). Second, migration of the CUB and TED domains is reflected by two subsets of cross-links. Five cross-links from TED to MG8, MG7, ANA and MG2 support the location of TED at “shoulder-height” as observed in the traditional view of the crystal structure of C3 (Fig. 5C) (13). In C3b, these cross-links were no longer detectable and had been effectively replaced by five TED-MG1 cross-links and a CUB-MG1 cross-link, locating the TED at the “bottom” of the β-chain key ring (14) structure (Fig. 5D), which is again consistent with the crystal structure. Third, distinct sets of cross-links for C3 versus C3b amongst the MG7, MG8, C345C and anchor domains suggest the rearrangements of domains in this region. Such a domain rearrangement is entirely consistent with a comparison of the crystal structures.

Interestingly, two pairs of residues, with one partner in TED and the other in MG1, were cross-linked in C3b, in solution, yet were far apart in the crystal structure (38.3 and 40.4 Å compared to a theoretical cross-linking limit of 27.4 Å for the cross-linker used here (Fig. 6A). A change in the juxtaposition of the TED and MG1 domain could explain this but, on the other hand, three further TED-MG1 cross-links support the arrangement seen in the C3b crystal structure. This apparent conflict can be resolved assuming the TED domain to be mobile with respect to MG1 in solution (Fig. 6B). This is consistent with several other indications that TED is mobile in C3b (28–30).

**Figure 6:**
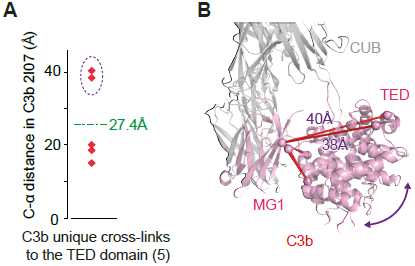
**Flexibility of the TED domain in C3b revealed by cross-linking data in solution. (A)** Two of five observed C3b-unique cross-links between the MG1 domain and the TED involve pairs of residues separated, in the crystal structure, by more than the theoretical maximum of 27.4 Å (Cα-Cα). (B) Putative flexibility of the TED domain in solution (indicated by the arrow) would explain the observation of these two over-length cross-links.

### QCLMS may reveal subtle protein conformational changes

Based on a comparison of QCLMS data and crystal structures, here we report some observations that may be more generally relevant to the challenging task of translating differences in yields of cross-linked products into inferred changes in protein conformation. Due to the absence of some residues in the crystal structures of C3 and C3b, not all cross-links can be evaluated against crystallographic evidence. Of the 104 cross-links observed, 71 bridged residues that are present in the crystal structures of C3 and C3b.

We first investigated the structural details of 25 cross-linked residue pairs that are unique to one of the two structures. As expected, six C3-unique and eight C3b-unique crosslinks agree only with the structure of the protein they were observed in but not the other when considering the residue pair distance in the crystal structures, offering an explanation for their absence based on distance solely. The remaining 11 C3/C3b-unique residue pairs, however, would be possible in both C3 and C3b when just considering the Euclidean residue pair distance in the crystal structures. To account for their absence from one or other structure, we must invoke steric effects that would prevent formation of a bridge (Fig. 7B, Supplemental Fig S1A-E), changes in accessibility of cross-linkable residues (Fig. 7C, Supplemental Fig S1F,G), or change of side chain orientation leading to increased distance of the reactive groups (Supplemental Fig S1I). For one case, a nearby sequence stretch absent from the crystal structure may have interfered with cross-linking (Supplemental Fig S1I).

**Figure 7:**
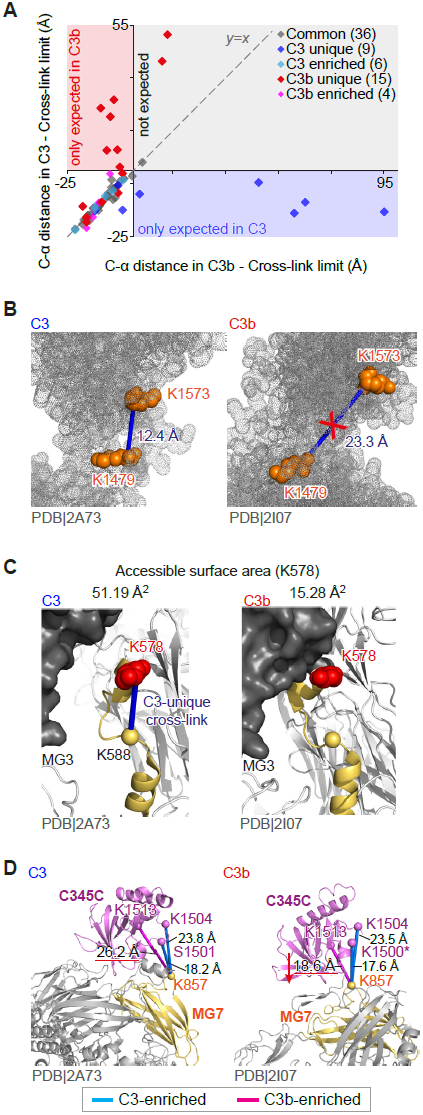
**Impacts of conformational changes on yields of cross-links. (A)** Measured distances (Cα-Cα) between cross-linked residues (in the C3 structure) minus maximal cross-linkable distance are plotted against the equivalents distances in the C3b structure (cross-links are only plotted where both cross-linked residues are present in the crystal structures). **(B)** Distance-wise, cross-link K1479-K1573 is within cross-link limit (27.4 Å) both in C3 (12.3 Å) and C3b (23.3 Å) structures. However these cross-links were only observed in C3 because, in C3b, the steric access of cross-linkers to both residues is blocked. **(C)** A dramatic decrease in the surface accessibility of K578 coincides with the absence of a C3 unique crosslink, K578-K588 (blue line), in C3b. Even though the proximities between K578 and K588 are nearly identical in the crystal structures of C3 (12.9 Å) and C3b (12.7 Å). **(D)** Decreased distance between residues K857^MG7^ and K1513^C345C^ may explain why their cross-link was significantly enriched in C3b. As an effect, other links involving K857^MG7^ namely K857^MG7^ – S1501^C345C^ and K857^MG7^– K1504^C345C^ are seen less in C3b. Residue K1501 is not present in the C3b crystal structure (PDB|2I07), residue 1500 is used instead for display purposes.

None of the cross-links that were quantified with ratios (nν46) involved dramatic proximity changes and the Euclidean distances between cross-linked residues in both crystal structures are within the limits of our cross-linker (Fig. 7A). However, we noticed that ten C3-enriched and C3b enriched cross-links exhibited bigger variations (1.8±2.3Å) on residue proximities between two crystal structures in comparison with those 36 mutual cross-links (0.8±1.1Å). In addition, C3-enriched and C3b enriched cross-links co-locate with C3-and C3b-unique cross-links by falling into the part of the molecule that experiences rearrangement during the transition from C3 to C3b (Fig. 5E). For seven of these ten cross-links, the crystal structures provided clues to explain the significant decrease on the yields of cross-linking in one structure versus the other: increase on residue distance (Fig. 7D), residue side chain orientation becoming less favorable for cross-linking (Supplemental Fig. S2A), change on residue flexibilities (Supplemental Fig. S2B), as well as influence from appearance or increase of colocated cross-links (Fig. 7D, Supplemental Fig S2C, D). The remaining three cross-links experience differences that cannot be rationalized by the crystal representations of C3 and C3b. However, cross-linking samples proteins in solution and the two proteins may differ in their insolution conformational ensembles. For this explanation also speaks that these three cross-links were found in the domains that experience rearrangements between C3 and C3b (Supplemental Fig. S2E).

## Discussion

We have established a workflow for QCLMS that we believe is in accordance with good practice in quantitative proteomics. This includes replication and label-swapping. In addition, we place the cross-linked pairs of amino acid residues at the focus of the analysis by gathering and summarizing all the relevant peptide quantitation data. We have lowered the barrier to entry for researchers wishing to apply this technique by establishing a semi-automated approach; this should also facilitate application of QCLMS to more complex systems involving, for example, multiple proteins. To achieve this, we enabled the standard proteomic quantitation software Pinpoint to work with data for cross-linked peptides by “linearizing” their sequences (Fig. 2A). Importantly, this mode of quantitation is independent of the specific algorithm used for identifying cross-linked peptides. As has recently become standard for all publications in *Molecular and Cellular Proteomics,* we propose that good practice in QCLMS would include open access to both the raw data and the lists of cross-links with associated quantitation. Our data are available via ProteomeXchange (25) with identifier PXD001675 and in the supplementary material.

Our study demonstrated that QCLMS is able to explore, in solution, the differences and similarities between the arrangements of domains in C3 and C3b. Cross-links that were unique or significantly enriched in one conformation over the other were observed in the parts of the molecules that experience major rearrangements between C3 and C3b. On the contrary, crosslinks that show no major changes on yield in C3 and C3b were detected in the parts of structure that are preserved from C3 to C3b. The excellent agreement of QCLMS-derived data with the structural transitions suggested by the crystal structures of C3 and C3b provide strong support for our approach. It also suggests some rules that determine how changes in protein conformation influence the yields of cross-linked peptides. Clearly, residue proximity can in many cases act as a simple binary switch for cross-link formation. But even in instances where a bridgeable distance between two cross-linkable residues is largely preserved during a conformational change (here as close as 0.1 Å), other factors than distance may impact crosslink formation; these include changes in surface accessibilities or the positions of the two partners relative to other structural features. While these factors could completely prohibit formation of a cross-link between two residues that are within range, complete negation seems to be rare. More commonly, these non-distance factors cause variation in yields of cross-linked products. This is manifested in a linkage that is enriched in one conformer relative to the other. In general, only complete loss of a cross-link may result from large conformational changes. In contrast, depletion or enrichment of the cross-link correlates with more local conformational changes that do not involve major distance change between cross-linked residues. It is therefore essential to distinguish experimentally between the two scenarios. This is possible through the robust quantitation, using replicated analysis with isotopic label-swapping, outlined herein

In conclusion, QCLMS is emerging as a tool for studying dynamic protein architectures. We have presented a carefully designed and thoroughly evaluated workflow for QCLMS analysis using isotope-labeled cross-linkers. A limitation of the approach is that each conformer to be analyzed must be stable on a time-scale during which it can be resolved from other conformers and then cross-linked. On the other hand, the automation in quantitation achieved in our protocols means that it is now feasible to extend a QCLMS study to the conformational changes that occur in multiple-protein assemblies (12). Thus it should be possible, for example, to follow the conformational preferences of a protein subunit in a series of assemblies as proteins are sequentially incorporated. The quantitation module in our workflow can also be adapted for SILAC-based quantitation or label-free quantitation (31) for cross-linked peptides. We envision that QCLMS will greatly facilitate the investigation of conformational dynamics in solution and help to animate the current largely crystallography-derived “snapshots” of biological processes.

## Acknowledgments

We thank Christoph Schmidt, Carla Clark and Paul McLaughlin for helpful discussions and Morten Rasmussen for initial software developments. We thank Sven Giese for helping providing annotated MS2 spectra. We acknowledge the PRIDE team for the deposition of our data to the ProteomeXchange Consortium. This work was supported by the Wellcome Trust (PNB: 081179, JR: 103139, Centre core grant: 092076, instrument grant: 108504) and BBSRC (BB/I007946/1). JR is a Senior Research Fellow of the Wellcome Trust.

